# Encoding Innately Recognized Odors via a Generalized Population Code

**DOI:** 10.1101/2020.01.28.923748

**Authors:** Qiang Qiu, Yunming Wu, Limei Ma, C. Ron Yu

## Abstract

Odors carrying intrinsic values often trigger instinctive aversive or attractive responses. It is not known how innate valence is encoded. An intuitive model suggests that the information is conveyed through specific channels in hardwired circuits along the olfactory pathway, insulated from influences of other odors, to trigger innate responses. Here we show that in mice, mixing innately aversive or attractive odors with a neutral odor, and surprisingly, mixing two odors with the same valence, abolish the innate behavioral responses. Recordings from the olfactory bulb indicate that odors are not masked at the level of peripheral activation and glomeruli independently encode components in the mixture. In contrast, crosstalk among the mitral/tufted cells changes their patterns of activity such that those elicited by the mixtures can no longer be linearly decoded as separate components. The changes in behavioral and mitral/tufted cell responses are associated with reduced activation of brain areas linked to odor preferences. Thus, crosstalk among odor channels at the earliest processing stage in the olfactory pathway leads to re-coding of odor identity to abolish valence associated with the odors. These results are inconsistent with insulated labeled lines and support a model of a common mechanism of odor recognition for both innate and learned valence associations.

## Introduction

Theories of sensory coding have always seen two major competing views, the specificity theory and the pattern theory (segregated vs. distributed processing). The specificity theory, often referred to as the “labeled-line”, suggests that sensory signals is processed along a fixed, direct-line communication to connect sensory input with a behavior [1]. The pattern theory, which has also taken terms like “across-fiber”, “parallel processing”, or simply “population code”, stipulates that information about a stimulus is distributed across different neurons and pathways. These theories have permeated discussions of sensory coding of all modalities since the mid-19th century. Whereas Adrian, von Frey, Barlow, Hubel and Wiesel, Mountcastle have all discussed the labeled line theory in their analysis of sensory responses [2], this type of circuitries mostly has been discovered for sensory inputs that carry innate valence information and trigger stereotypical responses. For example, the detections of looming threat and pain have been shown to be mediated by labeled line circuits [3-5]. In taste systems of insects and mammals, information about submodalities are found to be transmitted by separated pathways [6, 7]. In the fly olfactory system, there is strong evidence of labeled lines [8, 9]. Indeed, complete circuits that link sensory neurons expressing a specific receptor to the cells controlling behavioral output have been delineated [10].

The mammalian olfactory system is divided into at least four subsystems [11]. The vomeronasal organ, which is dedicated to detecting intra-species and inter-species cues [12, 13], is involved in eliciting innate, stereotypic behaviors and endocrine changes. Labeled lines have suggested to link specific pheromone cues to behaviors [14-16]. In the main olfactory systems, individual neurons express only a single receptor type and olfactory sensory neurons (OSNs) expressing the same receptor converge their axons into the same glomerulus in the olfactory bulb [17]. There is also a one-to-one connection between a glomerulus and the mitral/tufted cells. Despite the stereotypy in connectivity, studies have largely discovered a population code for odor identity, as individual odors generally activated multiple receptor types and each receptor can be tuned to many odors [18-20]. It seems that there is a dichotomy of coding paradigms that separates the vomeronasal system with labeled lines and the main olfactory systems using the population code.

However, behavioral and electrophysiological evidence has revealed innate connection between specific behaviors and individual chemosensory cues that activate the main olfactory system. Mice are attracted to food and urine of their own species, but they avoid scents from predators and odors associated with rotten food and decomposing carcasses. Single chemical compounds have been identified from predator urine, rotten foods, and decaying carcasses as signature odorants that directly elicit behavioral responses. For example, the compound (methylthio)-methanothiol (MTMT) has been shown as the active molecule that elicit attraction in male urine [21]. 2-methyl butyric acid (2-MBA), found in spoiled food, 2,3,5-trimethyl-3-thiazoline (TMT) from fox feces, and 2-phenylethylamine (PEA) from bobcat urine can directly elicit avoidance behaviors [22-26]. Similarly, the rabbit odor 2-methylbut-2-enal elicits suckling behavior in neonatal rabbits independent of other milk odors [27]. These odors are powerful drives of behaviors; the ability to properly recognize and respond to these odors is essential for survival of the animals in the wild. Importantly, these ethologically relevant odors have innately associated values, requiring no prior experience to elicit a response.

The innateness and the consistency of the response among different individuals, and the stereotypical response evoked by these odorants in different contexts suggest that inborn pathways transmit the information of these odors to evoke stereotypical behaviors. Indeed, it has been proposed that the activity of specific channels (receptors) convey valence information. For example, activation of Olfr1019, a receptor for TMT, induces immobility in mice, and knockout of Olfr1019 reduces the response [28]. Furthermore, it has been shown that Olfr288 mediates attraction by urinary odorant (Z)-5-tetradecen-1-ol (Z5–14:OH); low affinity activation of other receptors in the absence of Olfr288 leads to aversion [29]. PEA can activate the trace amine receptor TAAR4 at very low concentrations [30]. Knock out of TAAR4 abolishes low threshold aversive response to PEA, suggesting information about PEA may also pass through a highly specific pathway [24].

A variant of the specific channel hypothesis is that odor valence is topographically encoded, i.e., it is associated with specific regions of the olfactory bulb and in the brain. For example, MTMT activates the ventral, whereas TMT or 2-MBA preferentially activate the dorsal olfactory bulb [21, 22]. Genetic ablation of the dorsal olfactory bulb abolishes innate aversive responses to odors without affecting the learning of the same odors in general [22]. In whole brain single axon tracing of mitral/tufted cells, Igarashi and colleagues show that TMT and 2-MBA responding cells, which are located in the DII and DI zones of the olfactory bulb, respectively, project to the same region in posterolateral cortical amygdaloid area (PLCo) [22, 31]. Other studies have suggested that valence about aversive odors is transmitted through dedicated parallel pathways to ventral hypothalamus to elicit behavioral responses [32].

The notion of dedicated pathways transmitting innate valence information seems contradictory to the observation of a population code in the main olfactory system. Not only are individual mitral/tufted cells tuned to multiple odors [17, 33, 34], in areas traditionally thought to encode odor valence such as the PLCo, a distributed representation for odors with no apparent bias for the aversive odors are found [35]. For example, different ligands for the same TAAR can elicit opposing responses in a context-dependent manner [36]. Furthermore, odorants can interact to block attraction or aversion in a mixture without receptor antagonism [36]. These findings are consistent with the notion that most odor mixtures are perceived as configural, i.e., as a new odor, instead of as elemental (as individual components) [37]. However, it is difficult to reconcile a general population code with the innate nature of the responses. The ability for an animal to recognize the innately recognized odor independent of context also suggest that they are perceived as elemental, i.e., as separate components in the mixture. How these odors are encoded remain unknows.

In the labeled line model, innately recognized odors are expected to be encoded differently and separately from odors at large. The circuitry connecting the receptor neurons and brain centers that determine behavioral output is expected to be genetically hardwired and highly specific. However, without a clearly delineated circuits, it is difficult to address whether innately recognized odors are encoded by labeled line or by a general population code. All neural circuits require genetic specification to some extent, and population responses depend on the presence of specific set of neurons. Therefore, genetic intervention, optogenetic or chemicogenetic approaches designed to test the labeled line hypothesis can also alter population responses. For example, the loss of specific receptors may abolish the innate response to an odor, which can be interpreted as the loss of a labeled line. However, the same loss may generate an entirely different activity patterns and perceptual representation of odor identity. This interpretation would be consistent with a population code. Thus, the ambiguity in interpretations make it difficult to address how innate valence is encoded.

On the other hand, in a labeled line model one would expect the pathways transmitting odor information to be insulated from background and insensitive to environmental circumstances. Neural responses triggered by the innately recognized odor along the olfactory pathway are not expected to be altered by the presence of other odors. Thus, it is possible to examine whether mitral/tufted cells that encode innately recognized odors are subject to interference from other odors and whether activation of brain regions associated with innate perception and behavioral responses is altered. In this study, we take advantage of the assay using odor mixtures to examine behavioral responses to the innately recognized odors in the context of other odors. Our evidence points a model of innate odor coding using a generalized population code that is similar to those for learned odor preference.

## Results

### Odor mixing abolishes innate valence

In an odor-seeking assay, an animal’s approach toward the odor source is determined by at least three factors: novelty seeking, preference and odor habituation. For an attractive odor, prolonged investigation reflects attraction and novelty seeking. In contrast, an aversive odor may induce both novelty-seeking (for risk assessment) and avoidance. Odor preference is often assessed using place preference by associating odors with their spatial locations. However, intrinsic bias in spatial preference and scent marking can confound the readout [38, 39]. To avoid intrinsic place bias, we have devised a computerized setup, PROBES, to perform automated single-chamber odor preference assays [38]. This setup provides an efficient and robust readout of the sampling of a single odor port in the habituation/dishabituation test (Figure 1A). Using odors known to be attractive, neutral or aversive to mice, we examined odor investigation during first and second odor exposures after a period of habituation to clean air. The first exposure to mouse urines elicited vigorous investigation of the odor port, at a much higher level when compared with neutral and aversive odors (Figure 1B). Investigations of the aversive odors such as coyote urine, PEA were considerably less than those of neutral odors but similar to that of air (Figure 1C). When the mice were expose to the aversive odors a second time, investigation dropped significantly to below background level. The mice appeared to have avoided the odor port as exhibited by additional investigation once the odor delivery was stopped (Figure 1C). We reasoned that investigation during the first exposure likely reflected the effect of two competing drives: avoidance and risk assessment (investigation). During the second exposure, aversion likely became the dominant drive resulting in a decrease in investigation. To measure behavior preference, we used the average difference of investigative time between the first two odor epochs and the last air as a single index (Figure 1A). Attractive, neutral and aversive odor can be clearly identified with this index (Figure 1D).

**Figure 1.**
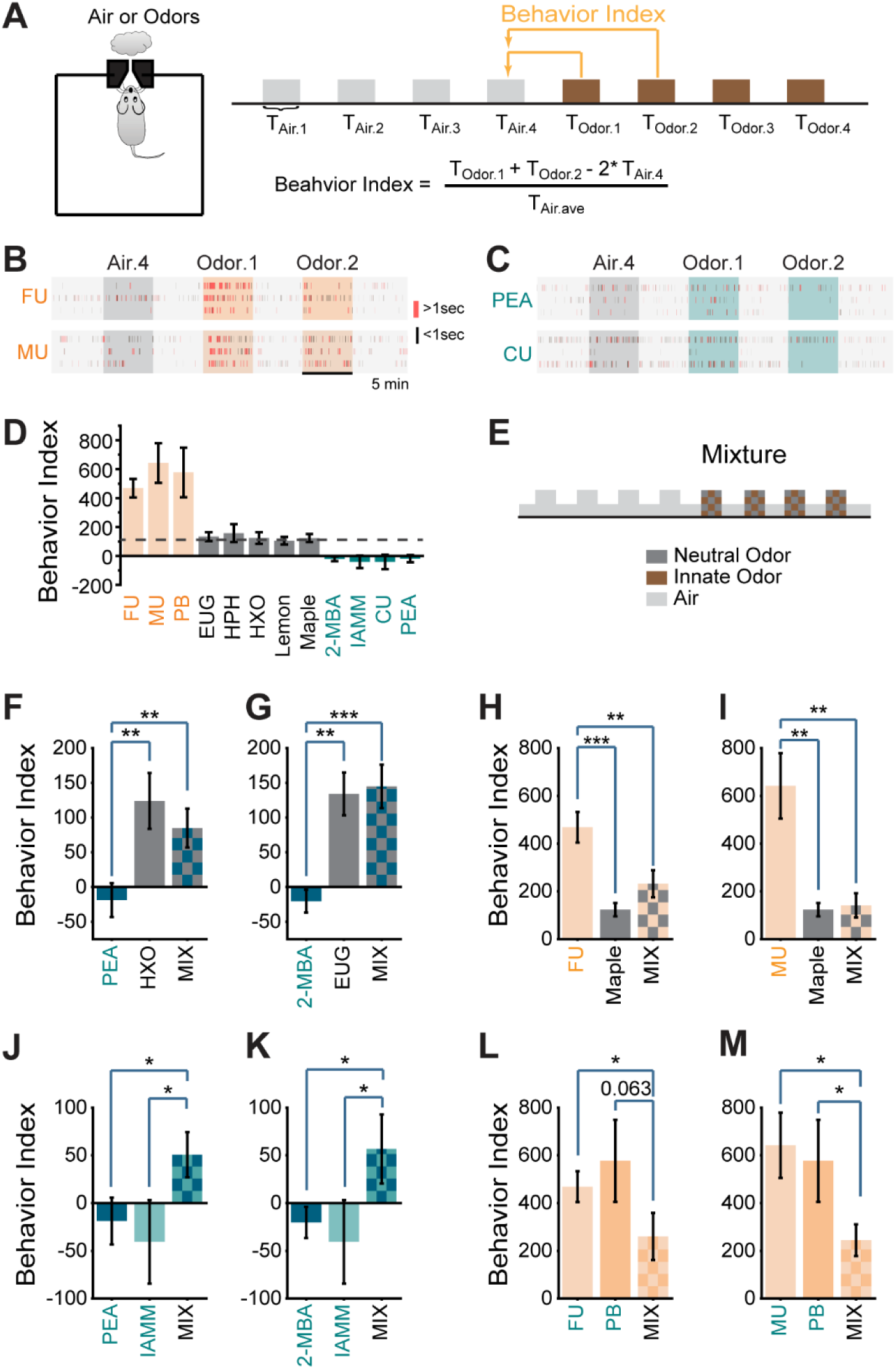
Odor mixtures abolish innate odor preference. (A) Illustration of the behavior paradigm. Left, single odor port arena. Right, odor presentation sequence and the quantification of behavior index. (B) Raster plots of odor port investigation (three animals each) of air, female or male urine (FU and MU) presentation. Only the fourth air presentation (Air.4, gray box) and the first two odor presentations (Odor.1 and Odor.2, orange boxes) are shown. Each tick represents an investigation event. Investigations longer or shorter than 1 second are marked by red and black ticks, respectively. (C) Same as (B) but for 2-phenylethylamine (PEA) and coyote urine (CU). Olive colored boxes indicating presentation of aversive odors. (D) Bar graphs showing behavior indices for a panel of odors (attractive: orange; neutral: gray; aversive: olive). Dashed line indicating the level for neutral odors. (E) Illustration of mixture experiment setup. (F, G) Bar plots of behavior indices measured for PEA, HXO and their mixture (MIX, F), and 2-MBA, EUG and their mixture (G). (H, I) Bar plots of behavior indices measured for female urine, maple and the mixture (H), and male urine, maple and their mixture (I). (J, K) Bar plots of average behavior indices to PEA, IAMM and their mixture (J), and 2-MBA, IAMM and their mixture (K). (L, M) Bar plots of average behavior indices to female urine, peanut butter and mixture (L), and male urine, peanut butter and their mixture (M). Dashed line indicating the level for neutral odors. One-way student *t*-test applied. * indicates *p* < 0.05, ** indicates *p* < 0.01, *** indicates *p* < 0.001, ns indicates *p* > 0.05. All bar graph data are shown in mean ± SEM.

We performed experiments using mixtures composed of equal parts of a neutral odor and an odor with innate valence (Figure 1E-I). Whereas PEA alone caused pronounced avoidance and 2-hexanone (HXO) did not, mixing PEA with HXO elicited no aversive response (Figure 1F). We observed similar effect for the mixture of 2-MBA (aversive) and eugenol (EUG, neutral; Figure 1G). A two-to four-fold change in the concentration of neutral odor in the mixtures did not affect this result (Figure S1). Similar observations were also made for attractive odors. Both male and female urine samples were attractive to mice of the opposite sex, but maple odor was not (Figure 1H). However, when maple was well mixed with the urine samples, the mixtures elicited little to no attraction (Figure 1H and 1I). In all pairs tested, odor preferences were abolished when innately recognized odors were mixed with neutral odors.

### Mixing aversive odors abolishes behavioral avoidance

How might mixing abolish innate odor preference? One possible scenario is that the presence of a neutral odor could overpower the aversive odor and mask its presence. Alternatively, the two odors may activate separate brain regions that drive competing behaviors, resulting in nullification of aversive responses. In both scenarios, if we mixed two odors of the same valence, the mixture is expected to elicit the same behavioral responses. We, therefore, performed odor preference tests using mixtures of PEA with 2-MBA, and of PEA with IAMM (Figure 1J and 1K). Strikingly, in both experiments, the mixtures did not elicit aversion. Similarly, well-mixed, innately attractive odors also reduced their attractiveness (Figure 1L and 1M).

### Linear decoding of odors from glomeruli responses

These behavioral experiment results were inconsistent with the labeled line hypothesis and suggest a different possibility. We hypothesized that mixing altered odor identities such that they were no longer recognizable. Volatile odors generally activate multiple receptors in the olfactory epithelium [18-20, 40]. Odor identity is encoded by the combinatorial activation of glomeruli, mitral/tufted cells in the olfactory bulb and by distributed sets of neurons in the cortices [41]. Recent studies indicated that there are interactions among odorants in activating receptors in mixtures [42, 43]. These interactions may create activity patterns that no longer be decoded as individual odors in the mixture.

We thus conducted a series of experiments to examine neural activities in the olfactory pathway. We first examined the activation of olfactory glomeruli by individual odors and their binary mixtures using the *OMP-IRES-tTA:tetO-GCaMP2* mice [12, 40]. Aversive odors including PEA, 2-MBA, IAMM, as well as neutral odors, including HXO and EUG, all activated the dorsal bulb. We presented EUG, 2-MBA and their mixture and recorded glomerular activation (Figure 2A and 2B). Mixtures elicited glomerular response patterns similar to the sum of the component odors. We plotted the response amplitude of the mixture against the arithmetic sum of the two individual odors. A simple linear relationship would indicate conformity between the two. The response amplitudes for individual glomeruli were mostly aligned with the diagonal line, indicating that the mixture did not cause overt masking at the level of the OSN that could explain the loss of aversion (Figure 2C). The same was observed for another pair of aversive/neutral odors, PEA and HXO (Figure S2A-C), as well as pairs of innately aversive odors including PEA/IAMM (Figure 2E-G) and PEA/2-MBA (Figure S2E-G). Even though some responses exhibited sub-linear summation for all the odor pairs we tested, the linear decoding from individual odors can predict the mixture response precisely using linear decoding of the patterns (Figures 2D, 2H, S2D and S2H).

**Figure 2.**
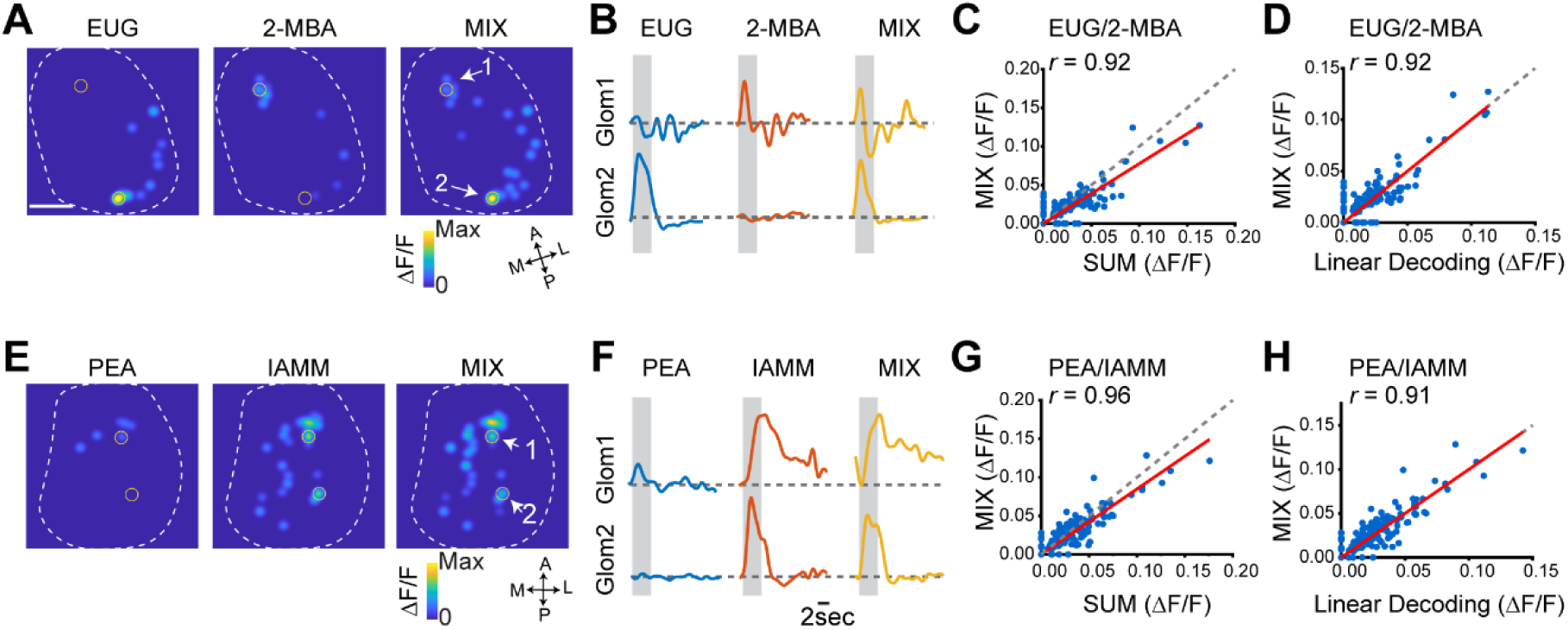
Linear decoding of odors from glomeruli responses. (A) Glomerular activation patterns elicited by EUG, 2-MBA and their mixture. Contours of the bulb are outlined. Orientations of the bulb are labeled as: A: anterior; P: posterior; M: medial; L: lateral. Scale bar, 500 µm. (B) Example response traces from the marked glomeruli in (A). Grey box indicates the odor presentation. (C) Scatter plot of glomerular response amplitude evoked by the mixture against the sum of response amplitudes to individual odors. Red line indicates linear fit of the data. Correlation coefficient (*r*) and diagonal line (grey dot) are indicated. (D) Scatter plot of response of individual glomeruli to the mixture against the predicted response from linear decoding for odor pair EUG/2-MBA. Red line indicates linear fit of the data. Correlation coefficient (*r*) and diagonal line (grey dot) are indicated. (E-H) Same as A-D, but for two aversive odor pair PEA/IAMM.

These observations showed that odor mixtures containing the innately recognized odors could be represented by linear combination of the glomerular responses to individual odors, similarly to other binary odor mixtures [44]. This result indicated that the glomerular representations of the odors were mostly independent of each other. If the insulated labeled line hypothesis is correct, this type of superposition would allow each odor to activate its distinctive pathway and elicit aversion.

### Altered odor representation by the mitral/tufted cells

Glomerular activities are transmitted to the mitral/tufted cells. These cells can sum input linearly in response to odor mixtures [45]. On the other hand, lateral excitation and inhibition mediated by interconnected neuronal networks may transform receptor activation into more complex population activities. To test whether the type of linear separation in the glomerular responses is carried through the olfactory pathways, we next examined neural activities of the mitral/tufted cells in the olfactory bulb. We performed two-photon calcium imaging experiments in *Cdhr1-Cre* mice injected with an adeno-associated virus (AAV) that expresses the calcium indicator GCaMP6f depending on Cre-mediated recombination (pAAV.Syn.Flex.GCaMP6f.WPRE.SV40) (Figure 3A) [46]. These mice expressed GCaMP6f specifically in the mitral/tufted cell population (Figure 3B).

**Figure 3.**
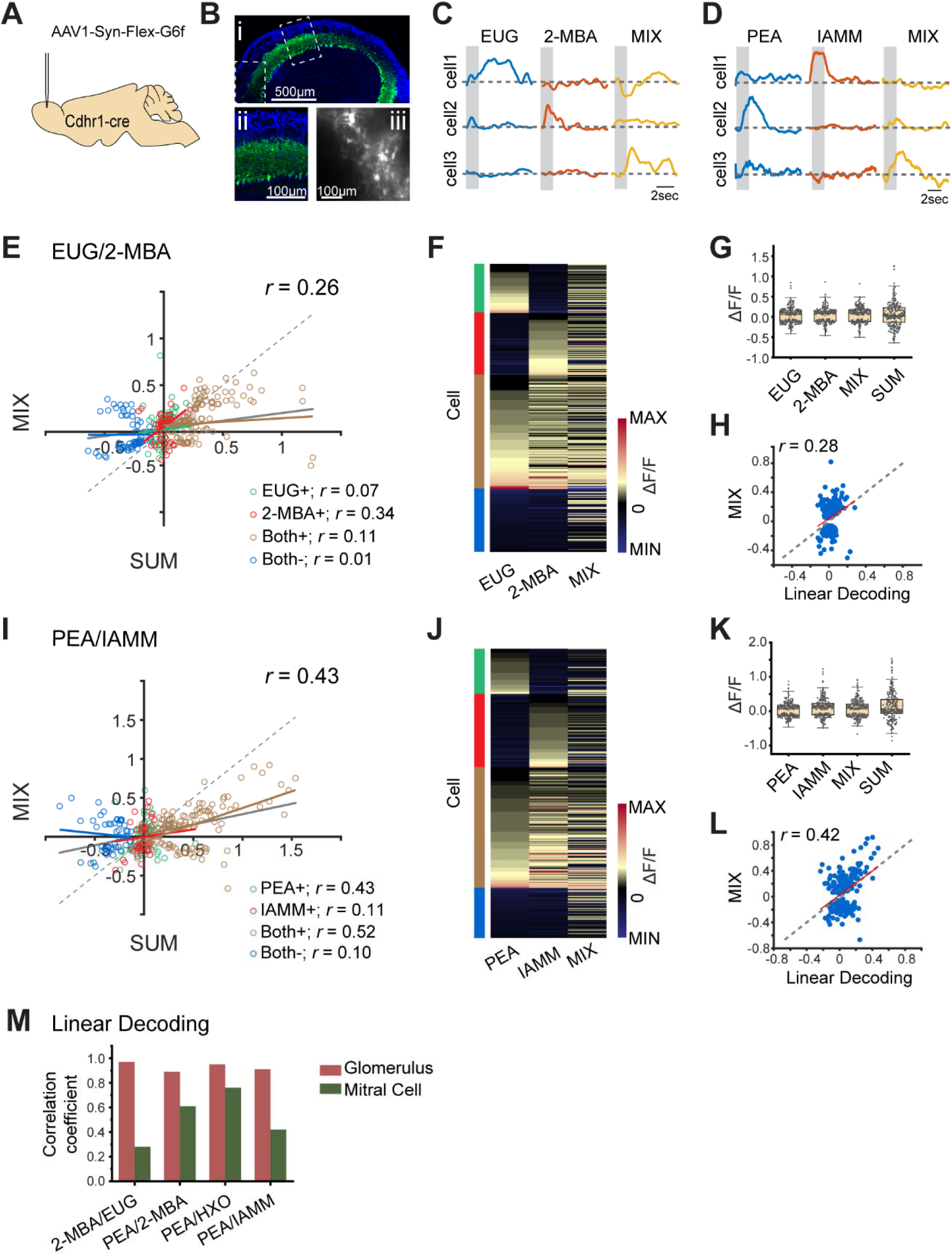
Cross talk among odor channels in the mitral/tufted cell population. (A) Illustration of the virus induced mitral cell GCaMP6f expression in mitral cells. (B) i, GCaMP6f expression in olfactory bulb after 3 weeks virus expression. ii, high magnification for the box area in i. iii, a 2-photon image showing the cells under recording condition. Green: GCaMP6f. Blue: DAPI. (C-D) Example response traces of mitral/tufted cell response to eugenol, 2-MBA and their mixture (C), or to PEA, IAMM and their mixture (D). Each row is a different cell. Gray box indicates odor delivery period. (E) Scatter plot of mitral/tufted cell response amplitude evoked by the mixture against the sum of response amplitudes to individual odors. Colored responses indicated positive response (increased calcium signal) to EUG only (EUG+), 2-MBA only (2-MBA+) or both odors (Both+), and negative response (decrease in calcium signal) to both odors (Both-). Gray line indicates the linear fit of all the data. Colored lines indicate linear fit of each group of data. Correlation coefficients (*r*) and diagonal line (grey dot) are indicated. (F) Heatmap shows mitral/tufted cell responses to EUG, 2-MBA and their mixture. Responses are sorted according to their response to EUG only, 2-MBA only, positive response to both odors and negative response to both odors. Groups are indicated with the colored bars that are matching the colors in (E). (G) Box plot shows the distribution of mitral/tufted cell response amplitude to odors: EUG, 2-MBA, their mixture and the arithmetic sum of component odors. Box plot edges indicate the first and third quartiles of the data, while whiskers indicate 1.5 interquartile range. (H) Scatter plot of response of individual mitral/tufted cells to the mixture against the predicted response from linear decoding for odor pair EUG/2-MBA. Red line indicates linear fit of the data. Correlation coefficient (*r*) and diagonal line (grey dot) are indicated. (I-L) same as (E-H) but for two aversive odor pair PEA/IAMM. (M) Bar graphs shows the correlations coefficients (*r*) between linearly decoded and the actual responses of the glomeruli (wine color) and mitral cells (olive color) for the four odor pairs.

We recorded mitral/tufted cell responses under awake conditions with precisely timed odor stimuli [38]. A total of 4320 odor/neuron pairs (288 cells and 15 odors) in response to PEA, IAMM, 2-MBA, EUG, HXO and their binary mixtures were recorded. The cells exhibited characteristic responses timed to odor stimulations with both increase and decrease in calcium signals (Figures 3C, 3D, S3A and S3B).

Quantitative measurements of the responses indicated that cells responding to only one of the two odors responded to the mixture with either enhanced or diminished amplitudes, and these bidirectional changes were observed for both aversive/neutral and aversive/aversive odor pairs (Figures 3E, 3F, 3I, 3J, S3C, S3D, S3G and S3H). For all odor pairs examined, response amplitudes of individual cells to the mixture were smaller overall than the arithmetic sum of responses to the components. This could be seen in pairwise plot of the response amplitudes for aversive/neutral (Figures 3E, 3F, S3C and S3D) and aversive/aversive pairs alike (Figures 3I, 3J, S3G and S3H). It could also be seen as the distribution of amplitude for the mixture, which was similar to those of single odors and smaller than the arithmetic sum (Figures 3G, 3K, S3E and S3I). This lack of summation as found in the glomerular response indicated that the overall response was normalized within the cell population.

The patterns of activity that represent the odor mixture were strikingly different from that for individual odors or their sum (Figures 3F, 3J, S3D and S3H). Using linear decoding from individual odor responses, we found that the mixture responses were poorly correlated with the predicted ones, indicating the mixture responses could not be linearly demixed into individual odor patterns (Figures 3H, 3L, 3M, S3F and S3J). Thus, the population response to the mixture likely represented a new odor identity rather than the combination of two components.

### Differential brain activation by aversive odor in mixture

The olfactory bulb projects to at least five cortical areas, including the posteriolateral cortical amygdala (PLCo) and the posterior nuclei of the medial amygdala (MeP), where the pathway appeared to diverge and activated different brain regions associated with different valence [47-50] (Figure 4A). Some of the brain regions activated along these pathways have been rigorously examined for their involvement of innate aversive or appetitive behaviors. For example, the ventral medial hypothalamic nucleus (VMH) mediates defensive behaviors including avoidance through activation of the anterior hypothalamic area, anterior part (AHA) [51, 52]. Projection from the ventral of posterior anterior medial amygdala (MePV) to the VMH has been shown to be involved in both attraction of mouse urine and aggressive responses [51, 53]. The medial amygdala (MeA), which receives input directly from the olfactory bulb, has been shown using optogenetic perturbation to be required to drive avoidance to predator odor through D1R-expressing cells projecting to the VMH [54]. MeA also projects to the bed nucleus of stria terminalis (BST), including the anteriomedial nucleus (BSTMA), to mediates anxiogenic response [55, 56]. Photo-stimulation of glutamatergic projection from BNST to the ventral tegmental area (VTA) triggers avoidance [57]. The AHA is required to trigger fear responses [55, 56, 58, 59]. Moreover, optogenetic intervention has been used to demonstrate that the PLCo mediates both innate aversive and appetitive behaviors in response to odors [60].

**Figure 4.**
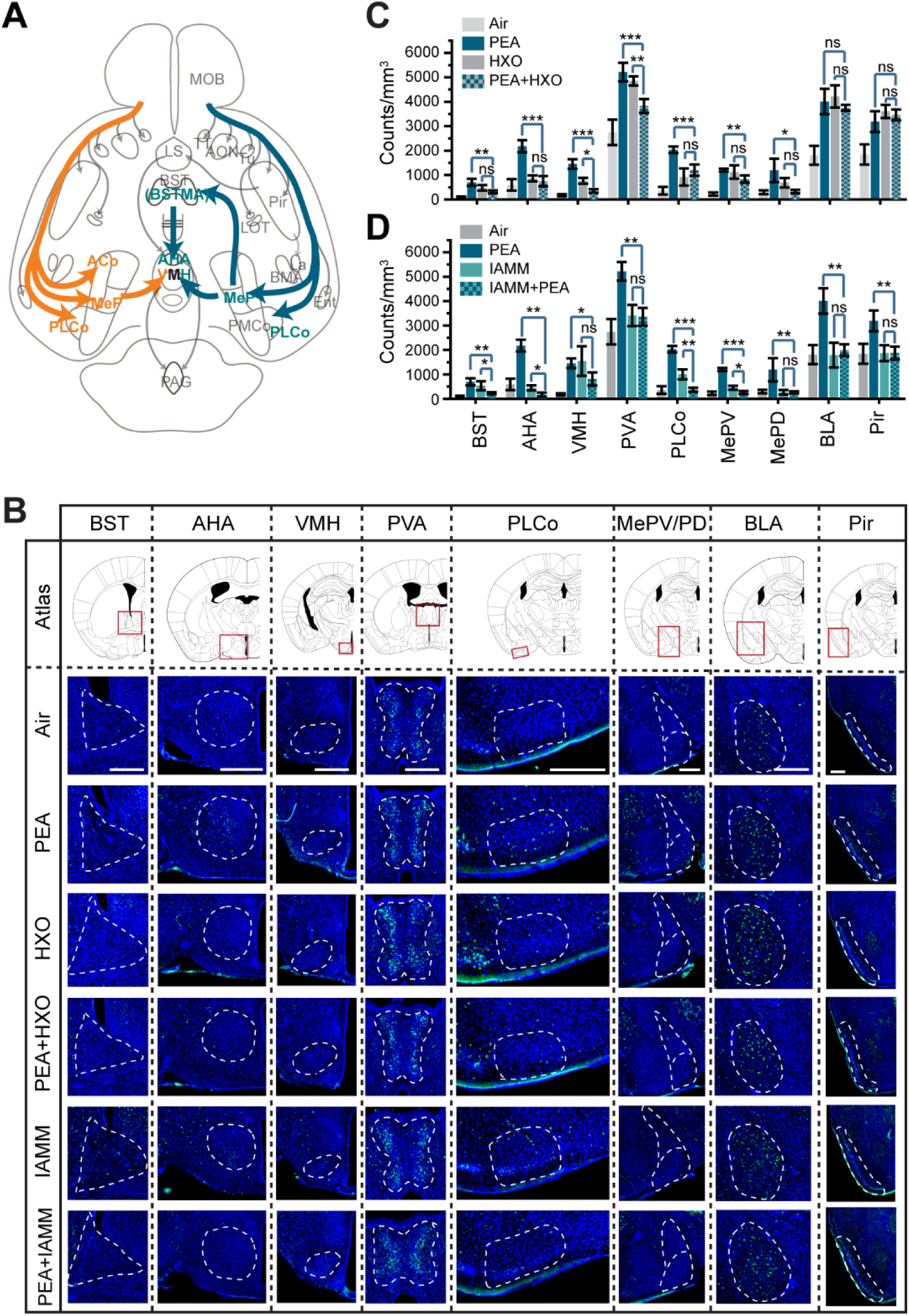
Altered representation of innate odors in the brain. (A) Illustration of pathways processing information about innately recognized odors Pathway and brain areas activated by attractive odors are marked by orange color. Dark olive color labels mark those for innately aversive odors. (B) Immunofluorescent staining of phospho-S6 (green) of brain sections from animals exposed to air, PEA, HXO, PEA+HXO, IAMM and PEA+IAMM. Cell nuclei were counter-stained with DAPI (blue). The areas being quantified are marked by dashed lines. Scale bars, 500 µm. (C) Bar plots of the density of activated cells in different brain areas for control, PEA, HXO and their mixture in B (data are shown in mean ± SEM, n = 6 half brains). (D) Bar plots show the density of activated cells in brain areas for control, PEA, IAMM and their mixture. Data are shown in mean ± SEM, n = 6 half brains. Abbreviation: AHA: anterior cortical amygdaloid area; AON: anterior olfactory nucleus; BLA: basolateral amygdaloid nucleus, anterior part; BMA: basomedial amygdaloid nucleus, anterior part; BSTMA: bed nucleus of the stria terminalis, medial division, anterior part; Ent: entorhinal cortex; La: lateral amygdaloid nucleus; LOT: lateral olfactory tract; MOB: LS: lateral septal nucleus; main olfactory bulb; MePD: medial amygdaloid nucleus, posterodorsal part; MePV: medial amygdaloid nucleus, posteroventral part; VMH: ventromedial hypothalamic nucleus; PAG: periaqueductal gray; PVA: paraventricular thalamic nucleus, anterior part; Pir: piriform cortex; PLCo: posterolateral cortical amygdaloid area; pmCo: posteromedial cortical amygdaloid nucleus; TT: tenia tecta; Tu: olfactory tubercle.

We sought to use the activation of these brain regions to further investigate odor coding. A dedicated channel that communicates the detection of a specific odor to these brain areas will lead to the activation of corresponding brain nuclei regardless of whether the odor is in a mixture. On the other hand, if the activation of these nuclei relies on the proper identity of the odor as encoded by the ensemble activity of mitral/tufted cells, then a change of the pattern of activity as we have recorded from the olfactory bulb would indicate that odor identities are altered in the mixtures and won’t activate the specific nuclei.

To quantify the activation of these brain areas, we performed immunofluorescence staining against phospho-S6 ribosomal protein (pS6), which stained activated neurons [61]. We sought to avoid influence of approach behavior on odor exposure by placing the odor vials in the home cage such as regardless of odor valence, the animals were exposed to the odors similarly. Following odor exposure, we performed serial sections through the brain to examine the patterns of activation in different regions and counted activated cells in 3-D volumes. PEA, but not the mixture with HXO, induced strong signals in BST, AHA, VMH, PVA, PLCo and MePV (Figure 4B and 4C). This observation suggested that areas associated with odor valence were differentially activated by an innately aversive odor depending on whether it was presented alone or in a mixture. The lack of activation by the mixtures was consistent with the lack of induced aversion and cannot be attributed to difference in approach behaviors. The number of cells activated in the basolateral amygdala (BLA) and the piriform cortex (Pir) were not significantly different (Figure 4B and 4C). Thus, areas that receive olfactory input, though not known to be particularly associated with valence, were similarly activated.

We then compared the brain activation patterns elicited by the mixture of two aversive odors with that activated by the aversive odors alone. Even though both PEA and IAMM activated the BST, VMH and PLCo, the PEA/IAMM mixture did not (Figure 4B and 4D). The lack of activation of these areas was consistent with the behavioral test results.

### Distinguishing aversive odors from background

It is counterintuitive that mixing an aversive odor with a neutral odor abolished aversive response. Animals react to predator or food odors in complex odiferous environments, suggesting that they can identify these odors in presence of different background odors. In our experiments involving odor mixtures, the two odors were presented contemporaneously. In natural environment, individual odors arrive at the nostril as plumes. The aversive odors could be detected with spatial-temporal displacement from environmental smells and be perceived as distinct. Thus, experimentally introducing temporal or spatial displacement in the presentation of the background odor may allow the aversive odors to be detected as elemental, i.e., as their own in the mixture. We tested this hypothesis by presenting one odor as background, followed by the mixture, and testing the innate response to the mixture (Figure 5A). In experiments using the neutral odor (HXO or eugenol) as background and testing with the mixtures with the aversive odor (PEA with HXO, or 2-MBA with eugenol), we found that the mixtures elicited the same level of aversion as the aversive odors alone (Figure 5B and 5C). In the converse experiments using the aversive odors as background, the mixtures were perceived as neutral (Figure 5B and 5C). Thus, background presentation of the neutral odors allowed PEA or 2-MBA within the mixtures to be properly recognized. Temporally displaced presentation of odors was sufficient to allow individual odors to be distinguished whereas well-mixed odors are not.

**Figure 5.**
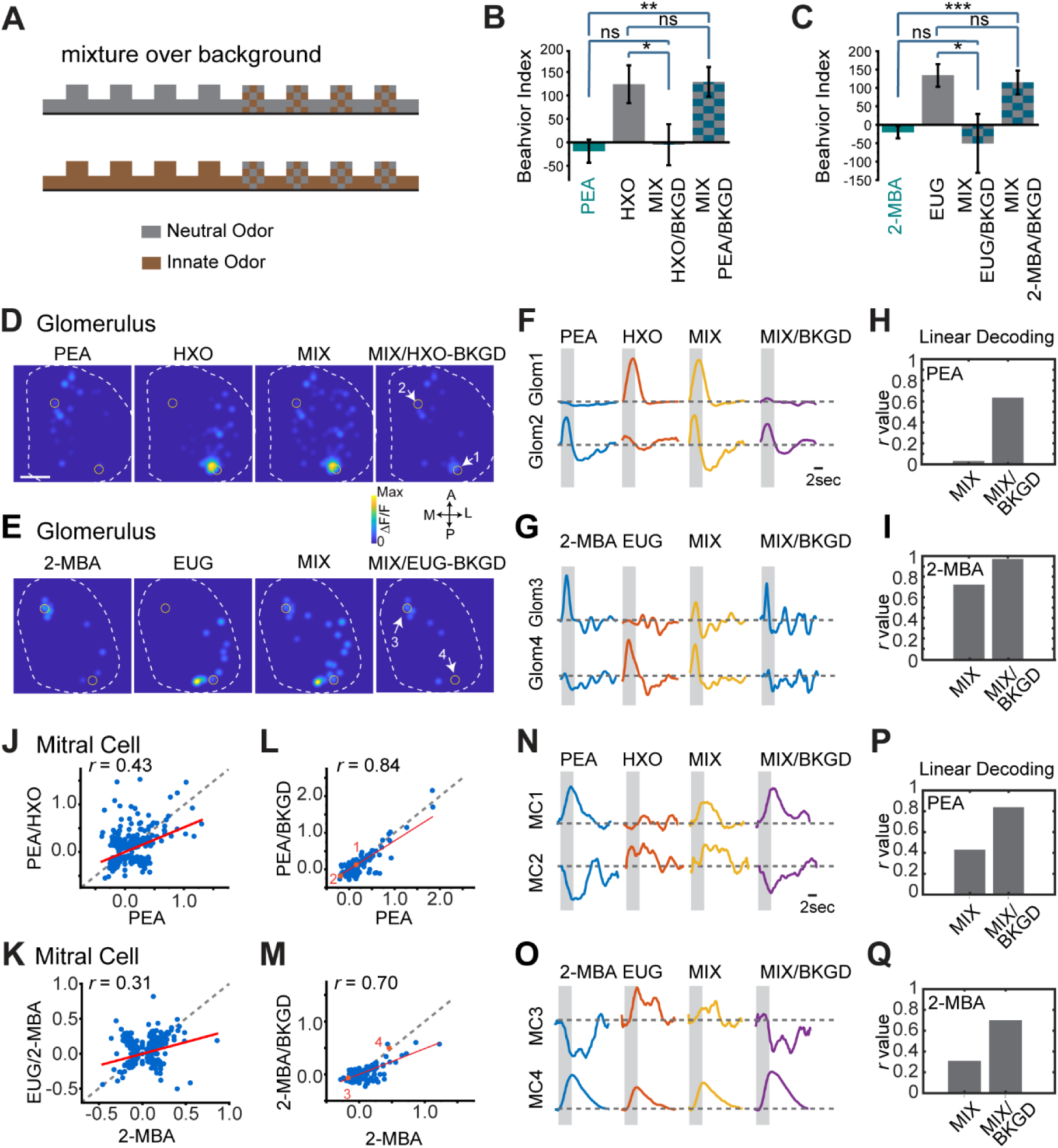
background presentation allows odor recognition. (A) Illustration of experimental setup. One of the odors (neutral or aversive) is delivered continuously as background. The mixture is delivered during the marked epochs. (B, C) Bar plots of behavior indices measured for PEA, HXO and the mixture when one of the component odors was presented as background (B), and 2-MBA, EUG and their mixtures when one of the component odors was presented as background (C). (D, E) Glomerulus activation patterns elicited by PEA/HXO (D) or 2-MBA/EUG (E) mixture after exposed to HXO (D) or EUG (E) as background. (F, G) Traces of glomerular (Glom, indicated in D and E) responses to individual odors, their mixture in the absence and presence of background presentation for odor pairs PEA/HXO (F) and 2-MBA/EUG (G) respectively. (H, I) Bar graphs shows the correlations coefficients (*r*) between linearly decoded mixture response and the actual glomerular responses to PEA (H) or 2-MBA (I) with or without background presentation. (J, K) Scatter plots of mitral cell responses to PEA/HXO mixture against that to PEA (J), and 2-MBA/EUG against 2-MBA (K) without background presentation. Red line indicates linear fit of the data. Correlation coefficient (*r*) and diagonal line (grey dot) are indicated. (L, M) Same as (J, K) but with the neutral odor presented as background. (N, O) Traces of mitral cells (MC, indicated in L and M) responded to individual odors, their mixture with or without background presentation for odor pairs PEA/HXO (N) and 2-MBA/EUG (O) respectively. (P, Q) Bar graphs shows the correlations coefficients (*r*) between linearly decoded mixture response and the actual mitral/tufted cell responses to PEA (P) or 2-MBA (Q) with or without background presentation. Correlation coefficient (*r*) values are plotted.

We next recorded glomerular responses to the mixture after presenting the neutral odor as background. We first recorded the responses to individual component odors and to the mixtures. The recording provided points for comparison. We then presented the neural odor as background, followed by the mixture, and recorded the response to the mixture. Following background presentation, the mixtures elicited responses that were dissimilar to those by mixtures alone. The overall patterns were similar to the innately aversive odors alone (Figure 5D-G). For individual glomeruli, the responses exhibited similar temporal dynamics and amplitude to that elicited by the aversive odors alone (Figure 5D-G). Linear decoding indicated that the mixtures elicited a response that can be predicted as the aversive odors (Figure 5H and 5I). Presenting the innately aversive odors as background, the mixtures elicited glomerulus responses that were similar to the neutral odors alone (Figure S4).

We also performed imaging of the mitral/tufted cells to monitor the response to the mixture after the neutral odor was presented as a background. Without background odor presentation, the mitral cell activity elicited by the mixture was poorly correlated with the pattern elicited by the aversive odor alone (Figure 5J and 5K). However, when the neutral odor was presented as background, the mixture elicited responses that were highly correlated with the aversive odors (Figure 5L and 5M). For individual cells, the dynamics and amplitude of the responses elicited by the mixture were similar to those elicited by the aversive odors (Figure 5N and 5O). Linear decoding indicating that mixtures presented over background were more likely linearly decoded as the aversive odors than the mixture alone (Figure 5P and 5Q). These adapted responses offered an explanation of the behavioral response to the mixture when presented over background.

## Discussion

Innate responses to some environmental stimuli are shaped by evolution to afford animals survival advantages even without learning. Because of the inborn nature of these responses, information about innately recognized cues is thought to be processed using circuits different from those for stimuli that can be learned by their association with unconditioned reward or punishment. Labeled lines, such as those mediating looming-induced flight response and spinal cord reflexes, provide a simple solution by linking sensory cells and behavioral centers (Figure 6A). Our results show that mixtures abolish innate aversive response. The most striking is that observation that mixing two odors with the same valence also abolishes innate responses. These observations cannot be simply explained by opposing actions of brain centers that drive different behaviors. Nor can it be accounted for by peripheral masking or antagonist interactions of ligands with the receptors [36]. Past studies have shown that presynaptic responses elicited by odor mixture were mostly mediated by intraglomerular interactions [44, 62, 63]. Consistent with these studies, glomerular responses to the mixtures can be linearly separated into those elicited by the component odors.

**Figure 6.**
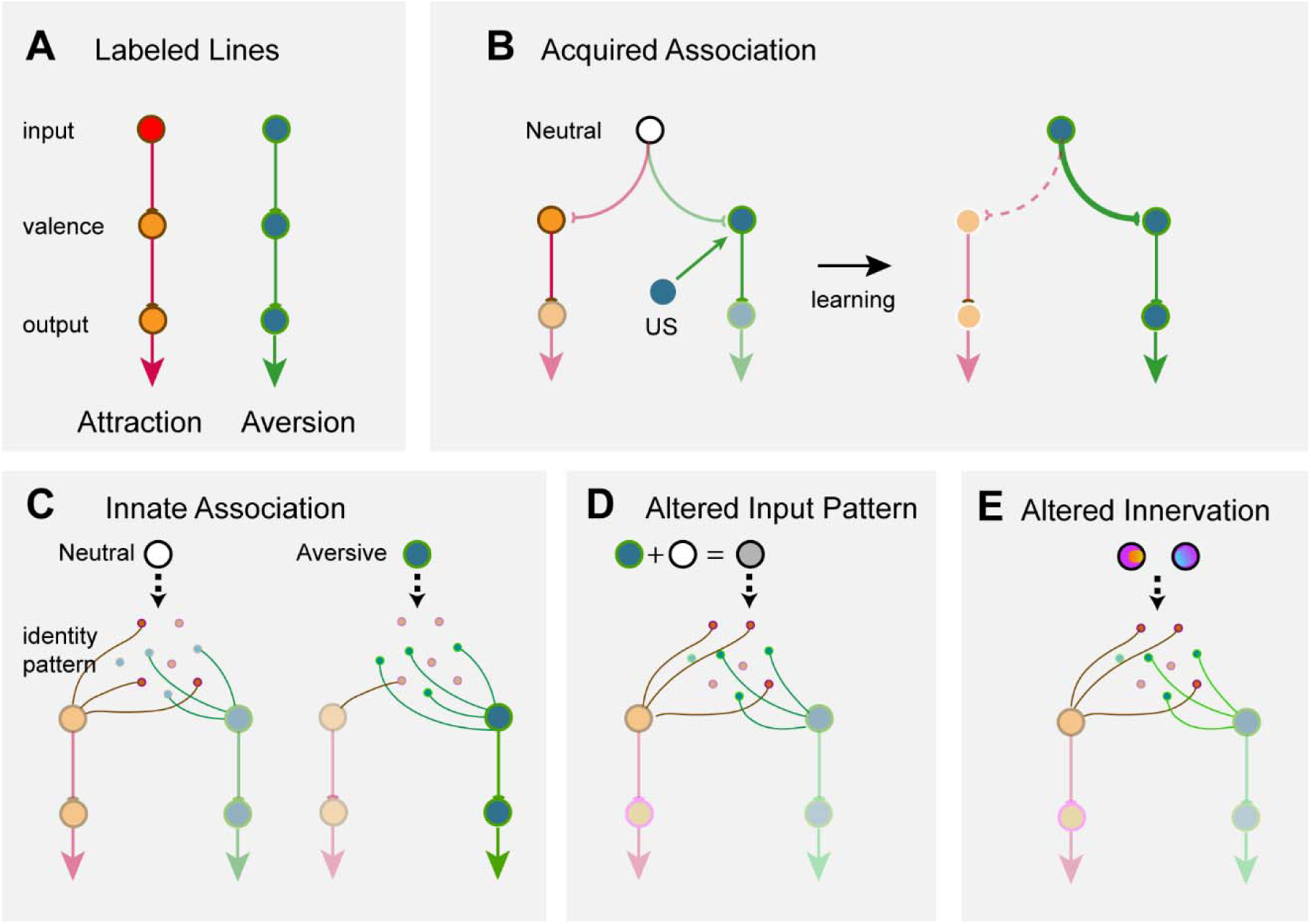
Models of encoding innate valence of odors. (A) The labeled-line model. Receptor activations are directly linked to behavioral outputs. (B) Acquiring valence through associative learning. A neutral stimulus has an unbiased connection to circuits leaning to attraction or aversion. A teaching signal such as an unconditioned stimulus (US) is associated with the neutral stimulus to enhance its connection to attraction, leading to learned response. (C) A model of innate odor preference. Neutral odors (white) activate sets of cells that do not have preferential connection to valence centers. Activation of a specific set of glomeruli (green) activates a set of cells, which encode the odor identity and are stereotypically connected to brain centers that assign valence. (D, E) Alterations in the activation of these cells, either through simultaneous activation of additional receptors (D) or through ectopic axon projections into multiple glomeruli (E), changes the identity of the odor being encoded and leading to changes in valence assignment.

A potential caveat is that our imaging study can only sample 10-15% of the dorsal olfactory bulb, a limitation on the experiment imposed by the anatomy of the rodent olfaction bulb. It is possible that the glomeruli imaged in this region exhibit linear summation but regions not imaged exhibit non-linear interactions. This is unlikely given the distributed nature of odor evoked response among the glomeruli. The likelihood that only the glomeruli out of the imaged area behave differently is small. In addition, the odor we tested activate the dorsal region more than the others. For example, 2-hexanone and eugenol have been shown by numerous studies using IEG staining and 2-DG uptake, to activate primarily the dorsal regions of the bulb [64-67]. PEA activates neurons expressing the TAAR4 receptors, which project to a few dorsal glomeruli [24, 30].

Importantly, there is a stark contrast between glomerular and mitral cell responses, both recorded from the dorsal bulb. Mixtures elicit responses in the mitral/tufted cell population that can no longer be linearly decoded. They do not conform to the individual channels that convey information of the component odors independently. The result does not fit the classic labeled line model where straightforward relays of information through the lines elicit stereotypical behaviors. Rather, they suggest that the highly interconnected networks in the mammalian olfactory pathway have a strong influence on how the innately recognized odors are encoded. This result differs from a previous study indicating that component mixtures can be linearly decoded using the mitral cell responses [68] and may reflect the differences in the odor set.

Natural odors are usually blend of individual odorant chemicals. Since individual chemicals activate multiple glomeruli and corresponding mitral cells, additive response from multiple chemicals in a blend would quickly saturate the responses. Presynaptic inhibition, which is mostly intraglomerular, as well as lateral inhibition among the mitral/tufted cells can reduce this saturation. Notably, interactions among the mitral/tufted cell population are mediated by inhibitory interneurons that are no long restricted to intraglomerular interactions [69]. The interglomerular interactions not only can normalize mitral cells response, they also can redistribute the patterns. The redistribution of mitral/tufted responses likely underlie the configural perception [37, 70] as the patterns elicited by the mixtures are treated as new rather than the combination of two separate known odors.

Innately recognized odors would be considered as elemental because it can be recognized in different odor environment. Notably, for many odors of this category, single molecules have been identified as innately aversive or attractive were from blends. PEA, TMT, MTMT and 2-MBA are all examples of single compounds identified from blend. However, our recorded mitral/tufted cell responses clearly show non-linear interactions in the mixture, which not only resulted in redistributed amplitude that appeared to have been normalized, but also resulted in patterns of activities that are likely perceived as novel to the animals. The results, therefore, suggest that the activity triggered by innately recognized odors are not insulated from those of others. In contrast, they are encoded similarly as all other odors, through a generalized population code resulting from interactions among the mitral/tufted cells. This redistributed activity, and the behavioral results, further suggest that mixtures containing these odors can nevertheless be perceived as configural.

How then, are these individual compounds perceived as independent of the context. We suggest the following possibilities. First, these compounds are more volatile. The volatility makes it possible, and likely, for these odor molecules to diffuse farther and separate from other molecules in the mix (the chromatography effect). Second, animals have evolved more sensitive receptors for these molecules, as exemplified by the extraordinary sensitivity to PEA and MTMT. The sensitivity would allow the single compounds to be detected before others, thereby create the temporal displacement needed for them to be perceived separately from other odorants. Alternatively, but not mutually exclusively, these compounds may be represented at a higher concentration in the source. Lastly, the receiving animals may have evolved a mechanism of feature selection, i.e., with specific detection of these molecules while filtering out activity evoked by other compounds [21]. These potential mechanisms may have created a situation where individual components in a mixture is detected as elemental, rather than configural.

If these innately recognized odors are encoded by a general population code, how are their valence assigned? Although the classic labeled line model (Figure 6A) provides a simple way of understanding intrinsic values associated, it is difficult to envision how it may operate with population activities that are subject to influence by other odors. Odor valence can be assigned through associative learning, presumably by associating the identify of an odor with an unconditioned stimulus (reward or punishment) to assign valence (Figure 6B). In the labeled line model, odor identity is associated with a specific channel, represented by a few highly specific cells insulated from interaction from other cells. Our results indicated that even for innately recognized odors, their identities are encoded by the ensemble activity of neurons in the olfactory pathway. We reason, therefore, that the encoding of innately recognized odor may be similar to those learned association (Figure 6C). In this model, odor identities are encoded by the population responses starting at the mitral/tufted cells and these identities are associated with valence. The difference is that for the innate odors this association is intrinsically determined rather than acquired through associative learning. This model allows genetic programs to specify the neuronal connection in a way that generates stereotypic patterns of activated cells to encode odor identities, and these cells have preset connection with valence circuit to drive approach or aversive behaviors. In contrast to the labeled lines, this model does not require neurons that encode innately recognized odors to be specifically tuned, nor does it require the activity encoding odor identities to be insulated from the influence of other odors. When the pattern of activity associated with an innately recognized odor is altered by mixing with another odor, the ensemble activity creates a novel odor identity and abolishes the valence associated with the component odor (Figure 6D). Activity-dependent modification of how olfactory sensory neurons project to the olfactory bulb during development would also alter these predisposed connections and change the valence associated with the odors (Figure 6E) [71].

This model can also explain the recognition of odors in the presence of background odors. When the odor presentation is temporally or spatially displaced from the background, individual odors in the mixture can be distinguished as distinct and their associated valence remain intact [72]. In this sense, innate detection of ethologically relevant odors is a form of elemental perception that requires their presentation to be independent from the environment. This model, therefore, allows a general coding strategy for all odors alike and permits flexible assignment of valence to neutral odors or reassigns valence to those with intrinsic preference.

## Methods

### Key Resources Table

| REAGENT or RESOURCE | SOURCE | IDENTIFIER |
| --- | --- | --- |
| Experimental Models: Organisms/Strains |  |  |
| Mouse: C57BL/6J | The Jackson Laboratory | JAX: 000664;<br>RRID:IMSR_JAX:000664 |
| Mouse: Cdhr1-Cre | MMRRC | 030952-UCD;<br>RRID:MMRRC_030952-UCD |
| Mouse: tetO-GCaMP2 | Yu Lab; The Jackson Laboratory | JAX: 017755;<br>RRID:IMSR_JAX:017755 |
| Mouse: OMP-IRES-tTA | The Jackson Laboratory | JAX: 017754;<br>RRID:IMSR_JAX:017754 |
| Odorants |  |  |
| 2-Hexanone | MilliporeSigma | CAS# 591-78-6;<br>Abbr: HXO |
| Heptanal | MilliporeSigma | CAS# 591-78-6;<br>Abbr: HPH |
| Eugenol | MilliporeSigma | CAS# 97-53-0;<br>Abbr: EUG |
| 2-Methylbutyric acid | MilliporeSigma | CAS# 116-53-0;<br>Abbr: 2-MBA |
| 2-phenylethylamine | MilliporeSigma | CAS# 64-04-0;<br>Abbr: PEA |
| isoamylamine | MilliporeSigma | CAS# 107-85-7;<br>Abbr: IAMM |
| Female mouse urine | MilliporeSigma | Strain: c57bl/6;<br>Abbr: FU |
| Male mouse urine | Fresh collected | Strain: c57bl/6;<br>Abbr: MU |
| Coyote urine | Harmon scents | Cat# CCHCY4;<br>Abbr: CU |
| Peanut butter | Jif Extra Crunchy | Abbr: PB |
| Lemon flavor | Frontier natural products co-op | Cat# 23071 |
| Maple flavor | Frontier natural products co-op | Cat# 23081 |
| Antibodies |  |  |
| Donkey anti-Rabbit 488 | Thermo Fisher Scientific | Cat# R37118;<br>RRID:AB_2556546 |
| Rabbit anti-Phospho-S6 (Ser235/236) | Cell Signaling | Cat# 4854;<br>RRID:AB_390782 |
| DAPI | Thermo Fisher Scientific | Cat# D1306;<br>RRID:AB_2629482 |
| Virus Strains |  |  |
| pAAV.Syn.Flex.GCaMP6f.WPRE.SV40 | Addgene | Cat# 100833-AAV1;<br>RRID:Addgene_100833 |
| Dental cement, Peptides and anesthetics |  |  |
| C&B Metabond Quick Adhesive Cement System | Parkell | Cat# UN1247 |
| 3-p peptide | United Peptide | biotin-QIAKRRRLpSpSLRApSTSKSE<br>SSQK |
| Isoflurane | Patterson Veterinary | Cat# 07-893-1389;<br>CAS# 26675-46-7; |
| Xylazine | Patterson Veterinary | Cat# 07-893-2121 |
| Ketamine | Vedco | Cat# VINV-KETA-0VED |
| Software and Algorithms |  |  |
| OriginPro | Origin Lab | https://www.originlab.com/Origin |
| MATLAB | Mathworks | https://www.mathworks.com |
| Custom MATLAB scripts | Yu Lab | N/A |
| ImageJ (Fiji) software | NIH | RRID:SCR_002285 |

### Lead Contact and Materials Availability

Further information and requests for resources and reagents should be directed to and will be fulfilled by the Lead Contact, Ron Yu.

### Experimental Model and Subject Details

#### Animals

The *OMP-IRES-tTA* (Jackson laboratory), *tetO-GCaMP2* (Jackson laboratory), and *Cdhr1-Cre* (MMRRC) mice were described previously [12, 40, 73]. The C57BL/6J (Jackson laboratory) were used for control group. All the animals were maintained in Lab Animal Services Facility of Stowers Institute at 12:12 hour reversed light cycle and provided with food and water *ad libitum*. All the behavior and functional imaging experiments were conducted during the dark light cycle under red or infrared light illumination. Experimental protocols involving mice were approved by the Institutional Animal Care and Use Committee at Stowers Institute and in compliance with the NIH Guide for Care and Use of Animals.

### Method Details

#### Odor delivery with olfactometer

Odor delivery was controlled by an automated olfactometer with custom written software developed in the National Instrument Labview programming environment [38]. Odorant chemicals are including: 2-Hexanone (Abbr: HXO), Heptanal (Abbr: HPH), Eugenol (Abbr: EUG), 2-Methylbutyric acid (Abbr: 2-MBA), 2-phenylethylamine (Abbr: PEA), isoamylamine (Abbr: IAMM) and were purchased from MilliporeSigma and freshly prepared in mineral oil at desired concentration. Female mouse urine (fresh collected from C57BL/6J mice, Abbr: FU), Male mouse urine (fresh collected from C57BL/6J mice, Abbr: MU), Coyote urine (Abbr: CU), Peanut butter (Jif Extra Crunchy, Abbr: PB), Maple flavor (Frontier natural products co-op, Abbr: Maple)), Lemon flavor (Frontier natural products co-op, Abbr: lemon) were used at original concentration. Odorants were then further diluted in carrier air with a maintained total flow rate (400 ml/min for calcium imaging and 100ml/min for behavior experiments).

#### Innate odor preference test

Experiments are the same as previously described [38]. 2-4-month-old mice were used for cross habituation experiments. Each experimental group contained 10-14 animals. Unless otherwise stated, all animals were naïve to the testing odors and exposed to the same odor once. Each animal was tested with a total of two separate experiments with at least one week between tests. After being habituated to the testing environment for half an hour, the animals were put into a 20cm x 20cm chamber for behavioral experiments. Odors from the olfactometer were delivered through a nose cone on one of the side walls, which is 5 cm above the base plate. A vacuum tube connected on the opposite wall of the nose cone provided an air flow to remove residual odors after odor delivery. Pure odorants were diluted into mineral oil at 1:10^3^ (v/v). The odors in the mixture were mixed at either the same concentration as individually (i.e., 100%A + 100%B), or at 0.5:1 ratio (50%A + 100%B), or at 0.25:1 ratio (25%A + 100%B). 10 ml/min air flow carried the saturated odor out from the odor vial and was further diluted into a 90 ml/min carrier air to make the final dilution to 10^−4^ (v/v). Delivery time, concentration and sequence of odor delivery were controlled by the olfactometer software. Investigation of odor source was registered by infrared beam breaking events and recorded by the same software that controlled the olfactometer.

The sequence of trial sessions is depicted in Figure 1. Odor was delivered for 5 minutes in each trial. After four trials of air presentation, a testing odor was presented 4 times. In a typical test, mice habituating to the test chambers over the multiple sessions of background air led to decreased *T*_*Air*_. The presentation of an odor elicited an increased *T*_*Odor*_. Repeated presentation of the same odor led to habituation, which was reversed by the test odor if it was perceived as novel. If the odor is attractive to the animal, an increased *T*_*Odor1*_ is expected as the mixed result of novelty and attraction. If the odor is avoided by the animal, a smaller increase or even decrease *T*_*Odor1*_ is expected as the mixed result of novelty seeking (risk assessment) and avoidance, while the *T*_*Odor2*_ is expected as the aversion only because the novelty is habituated quickly while the avoidance persists longer. To use the same index for both behaviors, we define the following behavior index:

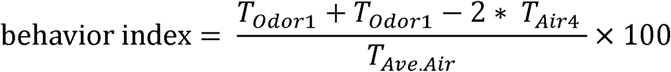

Odors used in innate behavior experiments were as follows: FU, MU, CU, PB, maple and lemon were delivered without dilution; PEA, IAMM, 2-MBA, HPH, HXO and EUG were diluted at 1:1000 in mineral oil. All the odors were then further diluted in the olfactometer in the air phase for another 10-fold.

### Phospho-S6 mapping of odor-evoked activity

For mixture phospho-S6 staining, animals were single housed and habituated in home cages for seven days with a glass vial covered with a plastic cover, which was punched with seven holes for odor evaporation while preventing physical contact with chemicals. For habituation, a small piece of cotton nestlet soaked with 500 µl mineral oil was put inside the vial. Vials were changed every day at one hour after light cycle. In day eight, a new glass vial with cotton nestlet soaked with 500 µl 2-MBA, PEA, IAMM, HXO or EUG at 1:10^3^ dilution in mineral oil, or freshly collected male or female urine was added in the home cages. Binary mixtures were prepared with equal mix of two odors. One hour after odor stimulation, mice were sacrificed and perfused with 4% PFA. The mouse brains were dissected and post-fixed with 4% PFA overnight at 4°C.

The phospho-S6 immunochemistry histology was performed based on the published protocol [61] with some modifications. The entire brain was cut into 50 µm thick serial sections using a Leica vibratome (VT1000S). Rabbit anti phospho-S6 antibody (Cell Signaling) was diluted 1:1000 in 0.1% PBST (0.1% Triton X-100 in 1X PBS) and used to incubate the slices at 4 °C overnight. The 3-p peptide (25 nM biotin-QIAKRRRLpSpSLRApSTSKSESSQK, where pS is phosphoserine; synthesized by United Peptide) was also included to reduce background staining [61]. Donkey anti-Rabbit 488 was (Thermo Fisher Scientific) diluted to 1:1000 in PBST for secondary antibody staining overnight. DAPI (Thermo Fisher Scientific) was used for nuclear staining.

Tiled images were acquired using Olympus VS120 Virtual Slide Microscope or PE Ultraview spinning disk confocal microscope (PerkinElmer), which were stitched together using the Volocity software (PerkinElmer). Different brain nuclei were identified based on the brain atlas (The Mouse Brain Stereotaxic Coordinates, third edition). The pS6 immuno-positive neurons were counted using ImageJ. For quantification, the two sides of the brain were treated independently and the following numbers of sections were used: anterior hypothalamic area, anterior part (AHA), 4 sections at -0.85∼-1.05 mm from Bregma; medial amygdaloid nucleus (posterodorsal area, MePD, and posteroventral area, MePV) and ventromedial hypothalamic nucleus (ventrolateral area; VMHvl), 5 sections between -1.35∼-1.60 mm from Bregma; hypothalamic nucleus (VMH) in the 2-MBA case, 4 sections between -1.80∼-2.00 mm from Bregma; bed nucleus of the stria terminalis (BST), 3 sections between 0.10∼0.25 mm from Bregma; posterolateral cortical amygdaloid area (PLCo), 5 sections at -1.35∼-1.60 mm from Bregma; paraventricular thalamic nucleus (PVA), 4 sections at -0.25∼-0.45 mm from Bregma ; piriform cortex (Pir), 5 sections at -1.35∼-1.60 mm from Bregma; basolateral amygdaloid nucleus, anterior part (BLA), 5 sections at -1.10∼-1.35 mm from Bregma.

### Awake head fixed calcium imaging

For calcium imaging of glomerulus, generation of the GCaMP2 mice was described previously [12]. Line 12i and 5i mice that exhibited strong fluorescence were used for imaging experiments as described previously [40]. One day prior to imaging, mice were anesthetized by intraperitoneal injection of ketamine/xylazine cocktail (100mg/kg, 10mg/kg body weight respectively) for surgery with thinned skull above the olfactory bulb.

For 2-photon mitral/tufted cell imaging, *Cdhr1-Cre* mice were anesthetized, and two small holes were made above each side of anterior and posterior olfactory bulb for virus injection. At each injection site, 250 nl pAAV.Syn.Flex.GCaMP6f.WPRE.SV40 virus (Addgene, 100833-AAV1, RRID:Addgene_100833) was injected into mitral cell layer at both 200 µm and 300 µm depths (Chen et al., 2013). After 3-4 weeks, animals were anesthetized to open a craniotomy window over the olfactory bulb and a 3 mm diameter glass cover slip was attached to the skull using dental cement. Animals were used for calcium imaging the next day.

Odorants were diluted in mineral oil (1:10^2^) and delivered at 40 ml/min flow rate. Total flow rate was maintained at 400 ml/min in all experiments. For the mixtures, both odors were delivered at 40 ml/min and mixed inside the olfactometer before delivered to the mouse nose. Odors were delivered for 3 seconds followed by a 27 second interval.

For GCaMP2 glomeruli imaging, responses to odor stimuli were collected on an Olympus BX60WI microscope using 4X air lens (Olympus XLFLUOR4X/340, NA 0.28). The image was collected at 512×512 resolution with 2×2 bin with sampling rate at 8.3 Hz. For mitral cell 2-photon imaging, the image was collected by an Olympus 2-photon microscope (FLUOVIEW FVMPE-RS) with 940 nm emission laser using 25X water lens (Olympus XLPLN25XWMP, NA 1.05). A resonate scanner with GaAsP detector was used for image collection at 512×512 resolution with 15 Hz sampling rate. Animals were head-fixed and freely standing on a roller during the experiment.

Custom-written scripts in ImageJ and MATLAB (Mathworks) were used for image processing as described previously [40]. Briefly, after ROIs were manually defined, the averaged response inside each ROI was extracted by ImageJ. Using customized MATLAB software, a baseline was defined for each ROI and ΔF/F was calculated. To display the response patterns, the peak amplitude of each glomerulus was mapped onto the spatial location of the glomerulus. The value was presented by applying a Gaussian blur with 50 µm standard deviation.

### Quantification and statistical analysis

All the statistics are conducted in MATLAB or OriginPro. Data were expressed as means ± SEMs in figures and text. Group differences were analyzed using one-way Student *t*-test. Significance was defined as: * indicates *p* < 0.05, ** indicates *p* < 0.01, *** indicates *p* < 0.001, ns indicates *p* > 0.05. MATLAB build in function ‘linsolve’ was used for the linear decoding analysis. It solves the linear system A*X = B using LU factorization with partial pivoting when A is square and QR factorization with column pivoting otherwise. In terms of the mixture case here, the formula is essentially the equation x1*A1+x2*A2=B. Here A1, A2 and B are the response vectors to the two component odors and the mixture, respectively. Each vector is the peak responses of N glomeruli (M/T cells). The linear system is expressed as matrix operation A*X=B, where A is a Nx2 matrix representing the component responses, B is a Nx1 vector, and X is 2×1 vector that is to be solved. A lower correlation coefficient indicates poor linear decoding.

## Acknowledgments

We thank A. Moran and members of the Lab Animal Services at the Stowers Institute for technical assistance. We also thank valuable input from members of the Yu laboratory. The work is supported by funding from Stowers Institute and the NIH (R01DC 008003, R01DC 014701 and R01DC016696).

## Declaration of Interests

The authors declare no competing interests.

## Supplemental Information

**Figure S1.**
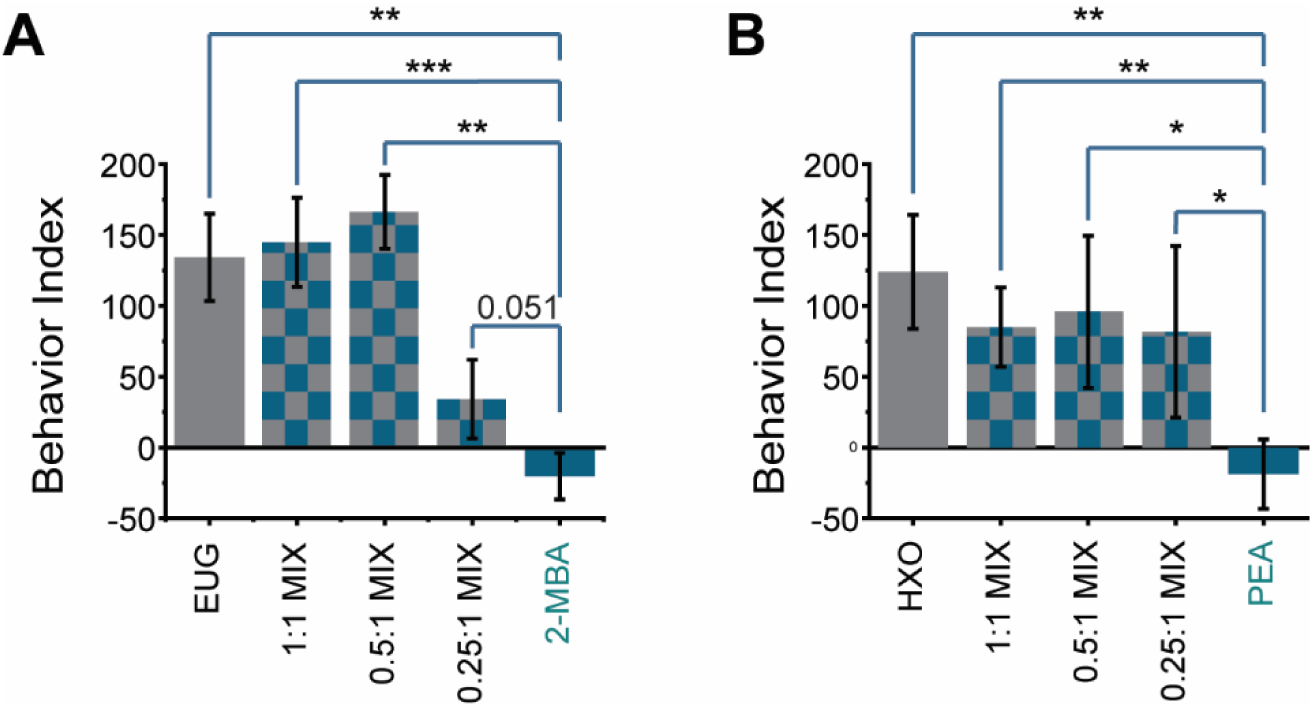
Odor mixtures with different mixture ratios abolish innate odor preference. (A) Bar plots of behavior indices measured for EUG, 2MBA and their mixtures with different ratios (1:1 mixture, 0.5:1 mixture and 0.25:1 mixture). (B) Bar plots of behavior indices measured for HXO, PEA and their mixtures with different ratios (1:1 mixture, 0.5:1 mixture and 0.25:1 mixture). One-way student *t*-test applied, * indicates *p* < 0.05, ** indicates *p* < 0.01, *** indicates *p* < 0.001. ns indicates *p* > 0.05. All bar graph data are shown in mean ± SEM.

**Figure S2.**
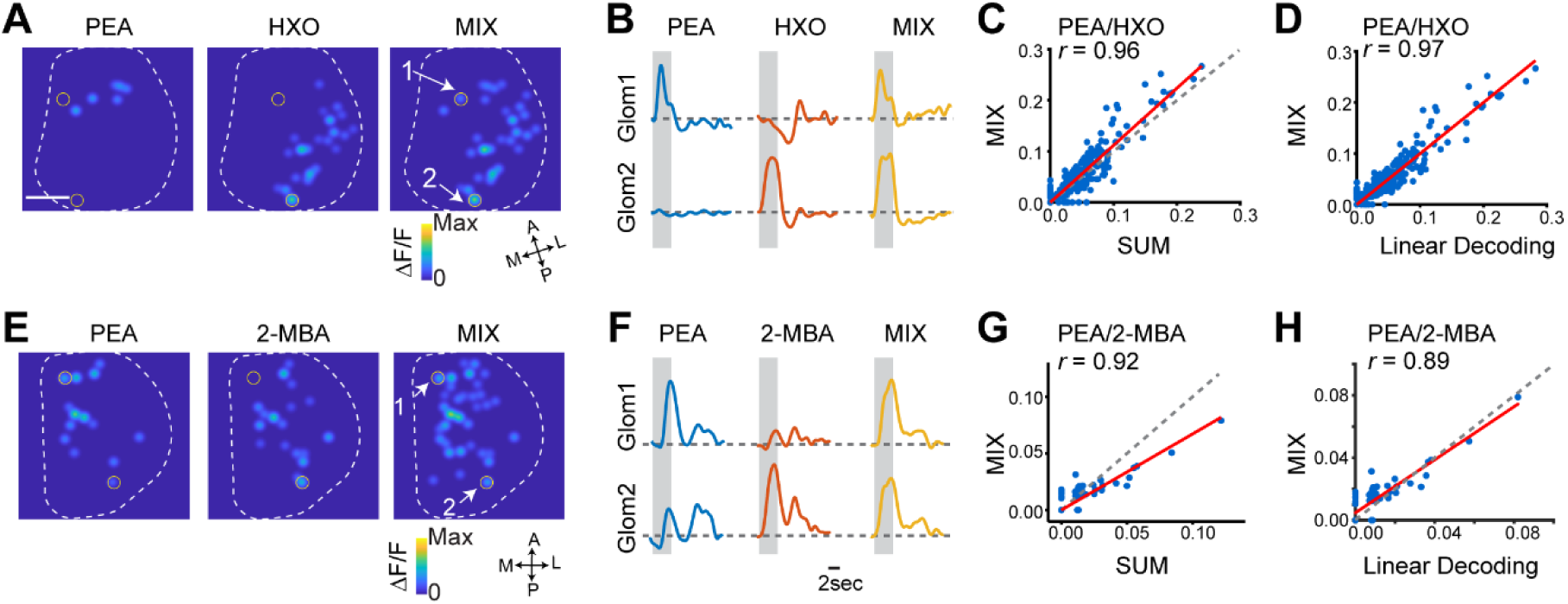
Linear decoding of odors from glomeruli responses. (A) Glomerular activation patterns elicited by PEA, HXO and their mixture. Contours of the bulb are outlined. Two of glomeruli are labeled. Orientations of the bulb are labeled as: A: anterior; P: posterior; M: medial; L: lateral. Scale bar, 500 µm. (B) Sample traces from the labeled glomeruli from (A). Grey box indicates the odor presentation. (C) Scatter plot of glomerular response amplitude evoked by the mixture against the sum of response amplitudes to individual odors. Red line indicates linear fit of the data. Correlation coefficient (*r*) and diagonal line (grey dot) are indicated. (D) Scatter plot of response of individual glomeruli to the mixture against the predicted response from linear decoding for odor pair PEA/HXO. Red line indicates linear fit of the data. Correlation coefficient (*r*) and diagonal line (grey dot) are indicated. (E-H) Same as A-D, but for two aversive odor pair PEA/2-MBA.

**Figure S3.**
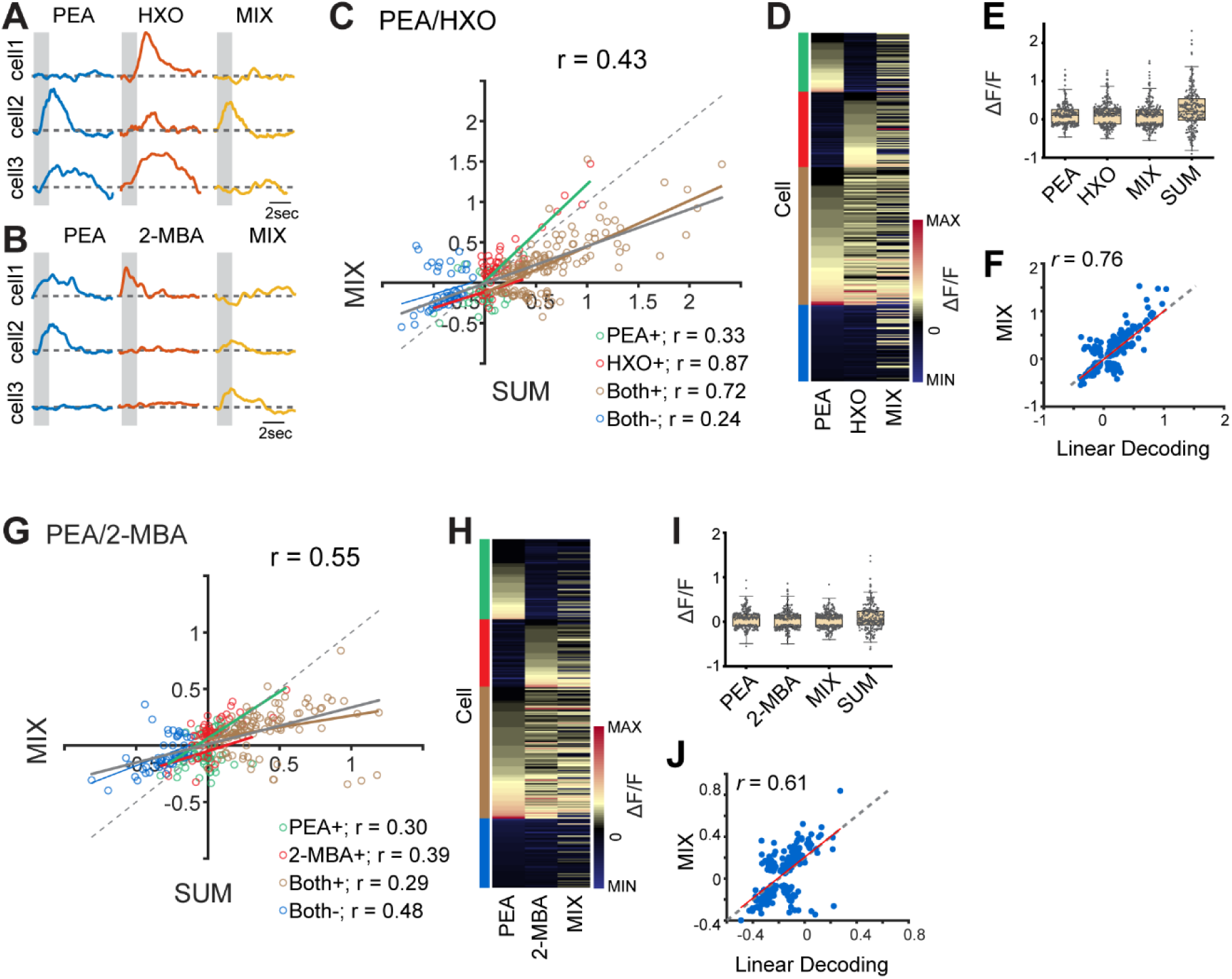
Cross talk among odor channels in the mitral/tufted cell population. (A-B) Example response traces of mitral/tufted cell response to individual odors PEA, HXO and their mixture (A), or to odors PEA, 2-MBA and their mixture (B). Each row is a different cell. Gray box indicates odor delivery period. (C) Scatter plot of mitral/tufted cell response amplitude evoked by the mixture against the sum of response amplitudes to individual odors. Colored responses indicated positive response to PEA only (PEA+), HXO only (HXO+), both odors (Both+) and negative response to both odors (Both-). Gray line indicates the linear fit of all the data. Colored lines indicate linear fit of each group of data. Correlation coefficients (*r*) and diagonal line (grey dot) are indicated. (D) Heatmap shows mitral/tufted cell responses to PEA, HXO and their mixture. Responses are sorted according to their response to PEA only, HXO only, positive response to both odors and negative response to both odors. Groups are indicated with the colored bars that are matching the colors in (B). (E) Box plot shows the distribution of mitral/tufted cell response amplitude to odors: PEA, HXO, their mixture and the arithmetic sum of component odors. Box plot edges indicate the first and third quartiles of the data, while whiskers indicate 1.5 interquartile range. (F) Scatter plot of response of individual mitral/tufted cells to the mixture against the predicted response from linear decoding for odor pair EUG/2-MBA. Red line indicates linear fit of the data. Correlation coefficient (*r*) and diagonal line (grey dot) are indicated. (G-J) same as (C-E) but for two aversive odor pair PEA/2-MBA.

**Figure S4.**
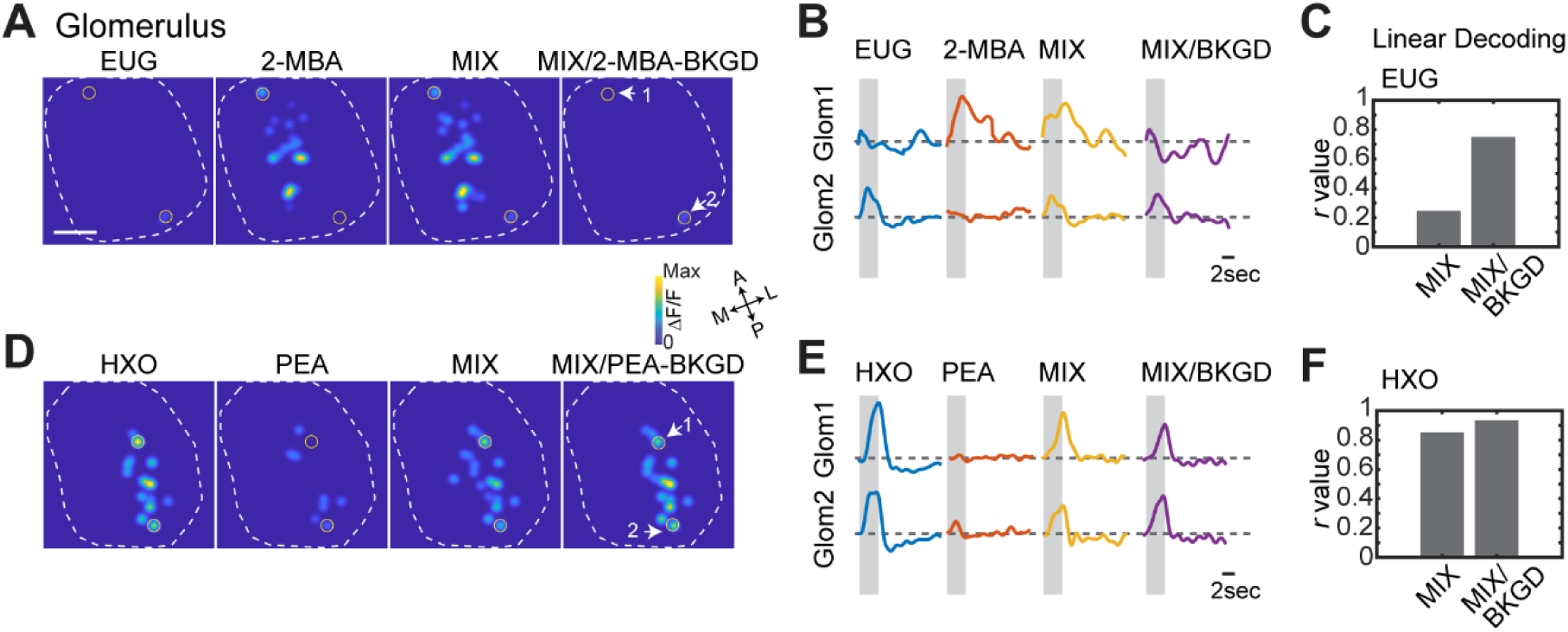
Glomerulus activity when presenting the innate odor as background. (A) Glomerulus activation patterns elicited by EUG/2-MBA mixture after exposed to 2-MBA as background. (B) Traces of glomerular (Glom, indicated in A) responses to individual odors, their mixture in the absence and presence of background presentation for odor pairs EUG/2-MBA. (C) Bar graph shows the correlations coefficients (*r*) between linearly decoded mixture response and the actual glomerular responses to EUG with or without background presentation. (D-F) Same as A-C, but for odor pair HXO/PEA with PEA as background odor.

## Notes

### Competing Interest Statement

The authors have declared no competing interest.

### Summary of Updates

Updated introduction, discussion, reference and a few new data added.

